# pH-responsive substrate switching in mycobacterial Type VII ESX secretion

**DOI:** 10.64898/2026.02.04.703728

**Authors:** Owen A. Collars, Richard L. Hernandez, Simon D. Weaver, Rebecca J. Prest, Caleb Manu, Gopinath Viswanathan, Rachel M. Cronin, Bradley S. Jones, David M. Tobin, Matthew M. Champion, Patricia A. Champion

## Abstract

During infection, pathogenic mycobacteria reside within phagosomes of varying acidity based on the macrophage activation state. The ESX-1 secretion system [early secreted antigen 6 kilodaltons (ESAT-6) system-1] delivers protein virulence factors essential for phagosome lysis, facilitating infection. The mechanisms underlying ESX-1 lytic activity in heterogeneous environments remain unknown. Here we show that the canonical Type VII secretion system, ESX-1, orchestrates substrate switching in response to different environments. Growing *Mycobacterium marinum* at acidic pH resulted in substrate switching *in vitro*. Substrate switching was accompanied by significant changes to the levels of ESX-1 substrate transcripts, and to the levels of both ESX-1 substrates and chaperones at the protein level. We showed that specific ESX-1 transcripts were significantly upregulated *in vivo*, and that distinct substrate sets are required in an acidic infection model.

**Significance Statement:** Pathogenic mycobacteria cause chronic and acute disease. Mycobacterial pathogens promote infection by transporting bacterial proteins into the host using ESX/Type VII secretion systems. The ESX-1 system secretes proteins into the phagosome that release the bacteria into the cytoplasm and promote bacterial survival in the macrophage. We show that *Mycobacterium marinum,* an animal pathogen and model for studying ESX-1 and tuberculosis, switches which ESX-1 proteins are secreted in response to acidic pH, an infection relevant signal. We demonstrate that protein secretion reflects changes in substrate transcripts and in substrate and chaperone protein levels. Finally, we leveraged two infection models to support that ESX-1 substrate switching likely occurs during infection. Our findings support a model in which mycobacterial pathogens use different proteins to lyse macrophage phagosomes of different pH.

## Introduction

*Mycobacterium tuberculosis*, the cause of human tuberculosis, transiently resides in the macrophage phagosome during infection (1–3). During this time, the bacteria sense the phagosomal pH and mediate a transcriptional response (4). The pH of the phagosome is dependent on the activation state of the macrophage (5). In naïve macrophages, mycobacteria arrest phagosome maturation, restricting the pH to a slightly acidic 6.0-6.5 (6). In activated macrophages, the phagosome fuses with the lysosome, relegating the mycobacteria to an acidic compartment with a pH of 4.5-5.0 (7).

ESX-1 is the canonical Type VII secretion system, and is essential for mycobacterial infection because it promotes phagosome escape (8–11). ESX-1 secretes protein substrates to the mycobacterial cell surface and into the environment (8, 12–17). Some ESX-1 substrates have been detected in the phagosome during macrophage infection (18) and several ESX-secreted proteins are presented as MHC-1 peptides (19). One or more ESX-1 substrates, along with virulence lipids, likely damage the phagosomal membrane, allowing mycobacterial pathogens access to the macrophage cytoplasm (11, 20–22). Cytoplasmic access activates host response pathways which result in macrophage lysis and bacterial spread (10, 23–27). It is unknown if the ESX-1 system differentially responds to changing environments during infection.

*Mycobacterium marinum,* a pathogen that causes a tuberculosis-like disease in poikilothermic animals, is an established model for studying *M. tuberculosis* infection biology (28–30). The ESX-1 system is functionally conserved between *M. marinum* and *M. tuberculosis* (31, 32). Importantly, *M. marinum* has been instrumental in dissecting the molecular function of the ESX-1 secretion system.

The precise machinery required for ESX-1 substrate translocation across the mycobacterial cell envelope is unknown. The ESX-1 substrates require each other for secretion (8, 16). Using proteomics and genetics, we proposed an “inside out” model for ESX-1 protein secretion in which secreted substrates span the mycobacterial cell envelope, promote the secretion of later substrates to the surface and out of the mycobacterial cell (Fig. 1). We identified four substrate groups [Groups I, II, III and IV,(33)]. The Group I substrates were required for the secretion of the other substrate groups. The Group I and Group II substrates were required for Group III substrate secretion. The Group II and Group III substrates were not required for Group IV substrate secretion. However, the secretion of Group IV substrates was higher from strains lacking the Group II or Group III substrates (33). We hypothesized that the Group II/III substrates and the Group IV substrates are secreted in opposition, potentially forming distinct substrate “assemblies” that function under different environmental conditions (Fig. 1). The different assemblies would require *M. marinum* to switch between secreting the Group II/III and the Group IV substrates. Although substrate switching is well defined in Type III secretion systems in Gram-negative bacteria, it has not been demonstrated for ESX secretion systems. For Type III systems, the destination of the secreted substrates acts as the signal for substrate switching. For example, in *Salmonella*, the neutral pH of the macrophage cytosol is the signal that promotes the switch from secretion system components to effectors (34). In this study, we leverage *M. marinum* to demonstrate ESX-1 substrate switching in response to physiologically-relevant pH changes during standard laboratory growth and during infection.

**Figure 1.**
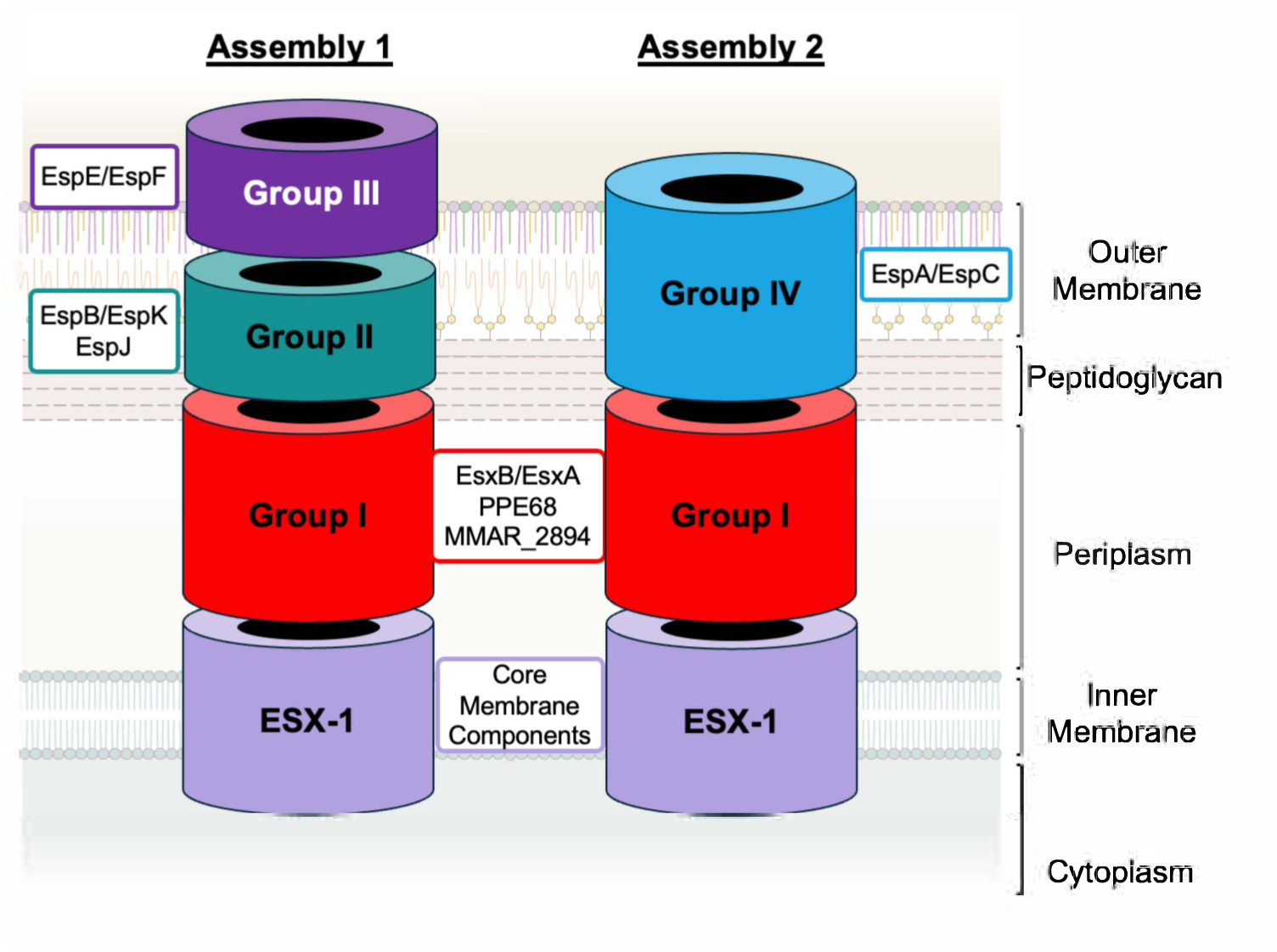
Model of proposed ESX-1 secretion Assemblies. Model based on proteo-genetic data from (33). Substrates and components drawn as rings for simplicity; based on reports of ring structures for EspB (86) and the ESX-1 membrane complex (87–90). Core membrane components include EccA-E(8, 91).

## Results

### The ESX-1 substrates are required for hemolytic activity change in response to environmental pH

The mycobacterial ESX-1 system secretes protein substrates into the macrophage phagosome (18). A unique aspect of the *M. marinum* model is that ESX-1 lytic activity can be measured *in vitro*. *M. marinum* lyse red blood cells in a contact-dependent, ESX-1-dependent manner (35, 36). Hemolytic activity simplifies measuring ESX-1 lytic activity because it does not require a host cell, and allows synchronization of lysis. We previously reported that deletion of individual substrate genes from Groups I, II or III significantly reduced or abrogated *M. marinum* hemolytic activity following growth under standard conditions. Hemolytic activity was restored by genetic complementation (17, 33, 37). Deletion of individual Group IV genes did not impact hemolytic activity (33).

To determine if hemolytic activity correlates with phagosomal lysis, we tested if individual ESX-1 substrates were required for phagosomal escape during macrophage infection. We previously adapted a luciferase reporter from *L. monocytogenes* that measures cytoplasmic bacteriolysis during macrophage infection (38, 39). Only *M. marinum* strains that lyse in the macrophage cytoplasm allow the reporter plasmid to be translocated to the nucleus where the luciferase genes are expressed. As shown in Figure S1A, a measurable population of *M. marinum* lysed in the macrophage cytoplasm resulting in luciferase detection, consistent with our prior report (38). The luciferase levels measured with the WT strain varied between infections, likely resulting from a lack of synchronized phagosome lysis. The Δ*eccCb_1_* strain lacks the ESX-1 membrane complex and is retained in the phagosome (8, 11, 40). Infection with the Δ*eccCb_1_* reporter strain resulted in a significant reduction in luciferase levels relative to infection by the WT reporter strain. We tested *M. marinum* strains lacking substrate genes representative from each group. The deletion of substrate genes in Groups I, II and III resulted in luciferase levels that were not significantly different from infection with the Δ*eccCb_1_* reporter strain (Fig. S1B-F). These data support that *M. marinum* strains lacking Group I, II or III substrates were retained in the phagosome. Complementation significantly increased luciferase levels relative to the Δ*eccCb_1_* strain in each case. Infection with the Δ*espC* (Group IV) reporter strain resulted in WT levels of luciferase that increased with complementation (Fig. S1G). These data support that the Group IV substrates are dispensable for phagosomal lysis under the conditions tested. Together, our data support that hemolytic activity correlates well with phagosomal lysis by *M. marinum*.

To determine if ESX-1 switches substrates, we tested if the requirements for specific substrates in ESX-1 mediated hemolytic activity changed in response to exposure to media buffered to different pHs. We reasoned that the requirements for hemolysis might change if substrate secretion switches in response to acidic pH. Using *M. marinum* strains with unmarked deletions of the known ESX-1 substrate genes (33), we exposed cultures to acidic (pH 5.0) or slightly acidic (pH 6.8) MOPS buffered 7H9 media for 24 hours (Fig. 2A). We washed the bacteria and measured the hemolytic activity of the *M. marinum* strains. As shown in Figure 2B, the WT *M. marinum* strain lysed RBCs following growth at pH 6.8 (filled circles) or pH 5.0 (open circles). Water and phosphate-buffered saline (PBS) were cell-free positive and negative controls for hemolysis, respectively. *M. marinum* strains with deletions in the membrane complex genes (*eccCb_1_,* light purple) or in the Group I substrate genes (*esxA, esxB, ppe68* or *MMAR_2894,* red) were non-hemolytic under both conditions tested. These data indicate that the ESX-1 membrane complex and the Group I substrates were essential for hemolytic activity following either growth condition. *M. marinum* strains with deletions of the genes encoding Group II (teal), or Group III (purple) ESX-1 substrates exhibited significantly increased hemolytic activity following growth at pH 5.0, compared to growth at pH 6.8 (Fig. 2B). These data support that Group II and Group III substrates were required for hemolysis under slightly acidic conditions, but dispensable at acidic conditions. *M. marinum* strains lacking the genes encoding the Group IV substrates (light blue) exhibited hemolytic activity following growth under either condition (Fig. 2B). The Δ*espC* strain was significantly more hemolytic following growth at pH 5.0 as compared to pH 6.8. These data support that the requirements for *M. marinum* hemolytic activity change in response to the pH of the growth media.

**Figure 2.**
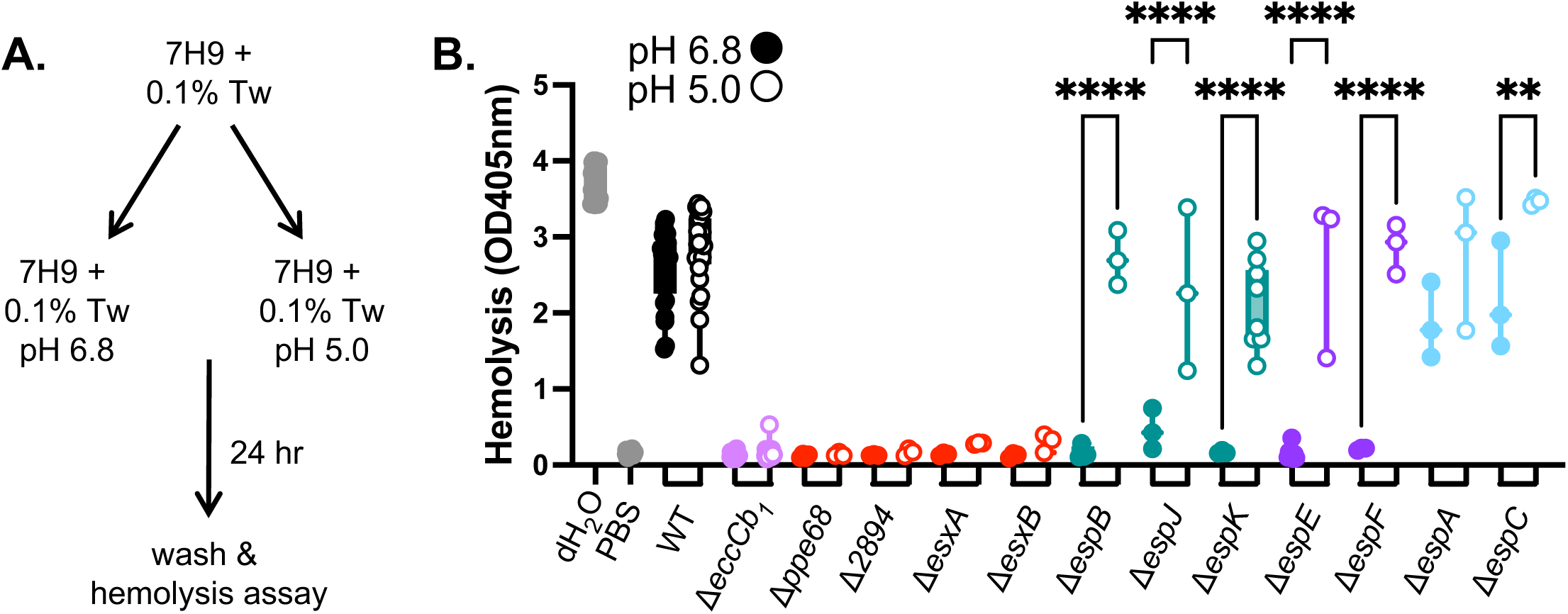
Exposure to acidic pH changes the requirements for hemolytic activity. (**A)** Workflow for acid stress exposure prior to hemolysis assay. Tw: Tween-80 **(B)** Hemolytic activity of *M. marinum* strains bearing single deletions at slightly acidic pH 6.8 (solid circles) or acidic pH 5 (open circles). Data points represent one biological replicate, which is the mean of three technical replicates. Significance was determined using a one-way ordinary ANOVA (*P*<0.0001) followed by a Tukey’s multiple comparison test. **** = *P*<0.0001, ** *P*=0.0023. Grey lines highlight the range of hemolytic activity from the WT strain. **** *P*<0.001. The double deletion strains were confirmed using PCR (Fig. S2), followed by targeted DNA sequencing.

### EspA and EspC are essential for hemolysis in the absence of Group III substrates

The data in Fig. 2 suggested that in the absence of the Group II/ III substrates, another group of ESX-1 substrates promoted hemolysis at pH 5.0 (Fig. 1). We generated a collection of *M. marinum* strains lacking the *espE* Group III substrate gene in combination with deletions of the other ESX-1 substrate genes (Table S1, Fig. S2). Deletion of the *espE* gene prevents the secretion of the other Group III protein, EspF, from *M. marinum* (33, 37). We tested the resulting double deletion strains for hemolytic activity following growth at pH 6.8 and pH 5.0. In agreement with Fig. 2B, the Δ*espE* strain was significantly more hemolytic following growth at pH 5.0 than at pH 6.8 (Fig. 3A). The hemolytic activity of the Δ*espE*Δ*esxA* and Δ*espE*Δ*esxB* strains (Group I, red) or Δ*espE*Δ*espA* and Δ*espE*Δ*espC* (Group IV, blue) were not significantly different following growth at pH 6.8 or pH 5.0. These data suggest that the Group I substrates, EsxA and EsxB, and the Group IV substrates, EspA and EspC, are required for the hemolytic activity of the Δ*espE* strain following growth at pH 5.0. Conversely, the Δ*espE*Δ*espJ* or Δ*espE*Δ*espK* strains (Group II, teal) were non-hemolytic following growth at pH 6.8, but were significantly more hemolytic following growth at pH 5.0, similar to the Δ*espE* strain. These data suggest that the Group II substrates, EspJ and EspK are dispensable for the hemolytic activity of the Δ*espE* strain when grown at pH 5.0. Together, these data suggest that the substrates required for the hemolytic activity of *M. marinum* switch in response to growth at acidic pH. Moreover, these data support that there are at least two sets or assemblies of ESX-1 substrates, Group II/III and Group IV, substrates that can promote the lytic activity of *M. marinum* (Fig. 1).

**Figure 3.**
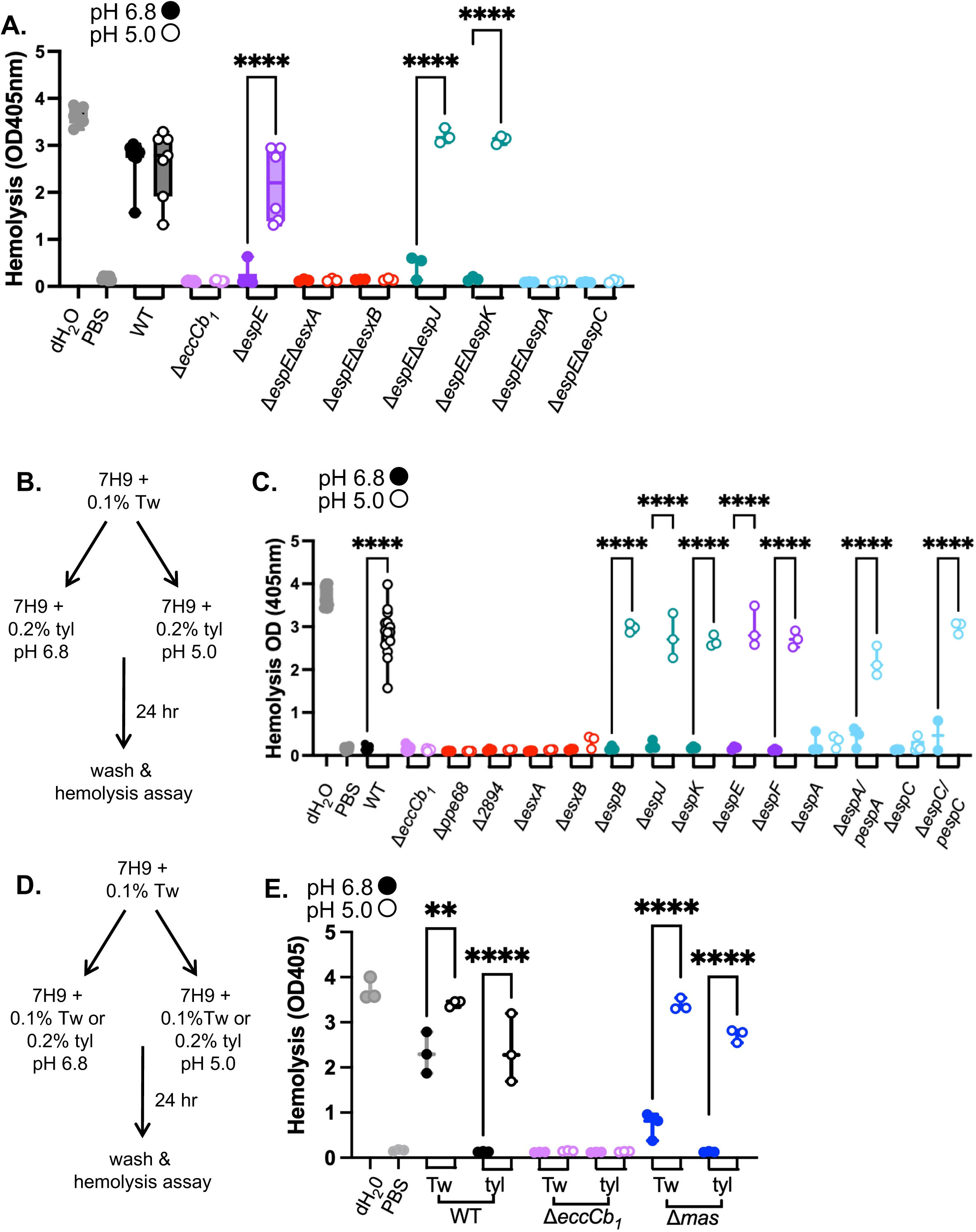
Group IV substrates are required for hemolysis in the absence of Group III substrates at pH 5. **(A)** Hemolytic activity of *M. marinum* strains lacking the *espE* gene (Group III) and representative substrate genes from each class following growth at pH 6.8 (closed circles) or pH 5.0 (open circles). Data points are individual biological replicates, which are the means of technical triplicate readings. Significance was determined using a one-way ordinary ANOVA (*P*<0.001) followed by a Tukey’s multiple comparison test. **(B)** Schematic of growth conditions for panel B., tyl = tyloxapol. **(C)** Hemolytic activity of *M. marinum* strains following exposure to tyloxapol and pH changes. Data points are individual biological replicates, which are the means of technical triplicate readings. Significance was determined using a one-way ordinary ANOVA (*P*<0.0001) followed by a Tukey’s multiple comparison test. **** *P*<0.0001. **(D)** Schematic of growth conditions for panel B., tyl = tyloxapol. Tw = Tween-80. **(E)** Hemolytic activity of *M. marinum* strains following exposure to tyloxapol or Tween-80 and pH changes. Data points are individual biological replicates, which are the means of technical triplicate readings. Significance was determined using a one-way ordinary ANOVA (*P*<0.0001) followed by a Tukey’s multiple comparison test. ** *P*=0.0027, **** *P*<0.0001.

In Type III secretion, substrate competition for the secretory machinery is an important aspect of substrate switching (41–45). The deletion of individual or multiple substrate genes could impact the mechanism of secretion or substrate selection. We sought to conditionally inhibit hemolytic activity without the genetic deletion of substrate genes to define which substrates were essential for hemolysis following growth under acidic conditions.

Based on our prior work, we reasoned that tyloxapol may be a chemical tool to block hemolysis mediated by Group II/Group III substrates, allowing us to test if the Group IV substrates mediate hemolysis in a wild-type genetic background (46). We grew the WT and Δ*eccCb_1_ M. marinum* strains, and those lacking individual substrate genes in 7H9 with 0.1% Tween-80 (Fig. 3B). We collected and washed the resulting *M. marinum* cells and resuspended them in 7H9 media buffered to either pH 6.8 or pH 5.0, with 0.2% tyloxapol for 24 hours. We washed the cells, and measured hemolytic activity. As shown in Fig. 3C, the WT strain, although non-hemolytic following growth at pH 6.8 (closed circles, consistent with (46)), exhibited significantly increased hemolysis following growth at pH 5 (open circles) in the presence of tyloxapol, similar to the cell-free water control. *M. marinum* strains lacking the *eccCb_1_* gene, the Group I substrate genes (*esxA, esxB, ppe68* or *MMAR_2894,* red) or the Group IV substrate genes (*espA, espC*, light blue) were non-hemolytic under either condition tested. Constitutive expression of the Group I genes (Fig. S3) or the Group IV genes in their respective deletion strains significantly restored hemolysis following growth in tyloxapol at pH 5, but not at pH 6.8. *M. marinum* strains lacking the Group II (*espB, espK* or *espJ,* teal) or Group III (*espE, espF*, purple) substrate genes were non-hemolytic following growth with tyloxapol at pH 6.8 but exhibited significantly higher hemolytic activity following growth with tyloxapol at pH 5.0. From these data, we conclude that the hemolytic activity of the WT strain following growth at pH 5.0 was independent of the Group II or Group III substrates. Moreover, we conclude that the EspA and EspC substrates can mediate hemolytic activity of *M. marinum* following growth at pH 5.0, but not at pH 6.8.

To rule out impacts of detergent or pH to PDIM (phthiocerol dimycocerosate)/PGL (phenolic glycolipid) virulence lipid levels (47, 48), which impact hemolytic activity (49, 50), we isolated total lipids from wild-type *M. marinum* grown in 7H9 media buffered to pH 6.8 or pH 5.0 with either 0.1% Tween-80 or 0.2% tyloxapol. We visualized PDIM, PGL and TAG (triacylglycerol) using thin layer chromatography. As shown in Fig. S4, all four conditions resulted in similar levels of detectable PDIM and PGLs. Growth at pH 5.0 with 0.1% Tween-80, and in the presence of 0.2% tyloxapol at either pH resulted in reduced TAG compared to the WT strain grown at pH 6.8 with 0.1% Tween-80. From these data we conclude that there were not gross changes to PDIM or PGL under the conditions used in this study.

We previously demonstrated that *M. marinum* strains lacking the PDIM and PGL virulence lipids exhibited reduced hemolytic activity, and that it specifically secreted significantly reduced levels of the EspE and EspF Group III substrates (50). We tested how the hemolytic activity of the Δ*mas* strain was impacted by growth in tween or tyloxapol, at pH 6.8 or 5.0 (Fig. 3D). As shown in Fig 3E, the Δ*mas* strain exhibited significantly reduced hemolytic activity compared to the WT strain following growth in media buffered to pH 6.8 with 0.1% Tween-80. The Δ*mas* strain was non-hemolytic following growth in media buffered to pH 6.8 with 0.2% tyloxapol, supporting that EspE and EspF mediate the reduced hemolytic activity in this strain. Upon shifting to growth in media buffered to pH 5.0, the Δ*mas* strain was significantly more hemolytic regardless of growth in either detergent. These data support that substrates other than EspE and EspF can promote the hemolytic activity of *M. marinum* following exposure to acidic conditions.

### *M. marinum* switches substrate secretion in response to acidic pH

We established conditions to measure ESX-1 substrate production and secretion from *M. marinum* in response to the slightly acidic (pH 6.8) or acidic (pH 5.0) conditions used in our hemolysis assays. We grew *M. marinum* in 7H9, and then diluted to an OD_600_ of 0.8 into 7H9 media buffered to pH 6.8 or pH 5.0, or Sauton’s defined media for 48 hours, and tested for the secretion of the Group I substrate, EsxB. EsxB (CFP-10) is one of the canonical ESX-1 substrates used to optimize the secretion assay in Sauton’s media in the initial ESX-1 secretion papers in *M. tuberculosis, M. smegmatis* and *M. marinum* (8, 36, 51). As shown in Figure S5, EsxB was secreted from the WT strain under all three conditions, although to a lesser extent from the WT strain grown in the 7H9 media buffered to either pH (lanes 1 and 2) as compared to Sauton’s media (our standard protocol, lane 3). Deletion of the *eccCb_1_* gene resulted in a loss of EsxB secretion under all conditions tested (lanes 4-6).

We reasoned that the differential requirements for hemolysis may reflect a switch between which ESX-1 substrate secretion in response to acidic pH. Our data in Fig. 3 suggests that both Group III (EspE/F) and Group IV (EspA/C) substrates promote hemolysis following growth at pH 6.8, while the Group IV substrates (EspA/EspC) only contribute to hemolysis following growth at pH 5.0. We hypothesized that the secretion of the Group III and Group IV substrates switch in response to the pH. We generated cell-associated and secreted protein fractions (Fig. 4A) and detected EspE and EspA production and secretion using immunoblot analysis. As shown in Fig. 4B, the EspE protein (purple arrow) was produced in the WT and Δ*eccCb_1_* strains during growth in media buffered to pH 6.8 (lanes 1 and 3), and was secreted from the WT strain (lane 5) but not the Δ*eccCb_1_* (lane 7) strain. While EspE was detected in the cell-associated fractions of the WT and the Δ*eccCb_1_* strains grown at pH 5.0 (lanes 2 and 4), the levels of EspE were reduced. The EspE protein was not detected in the secreted protein fractions of the WT or the Δ*eccCb_1_* strain following growth at pH 5.0. In contrast, the EspA protein (light blue asterisk) was detected in the cell-associated protein fractions from the WT and Δ*eccCb_1_* strains following growth in media buffered to pH 5.0 (lanes 2 and 4), but not pH 6.8 (lanes 1 and 3). EspA was detected in the secreted fraction generated from the WT strain following growth at pH 5.0 (lane 6). From these data, we conclude that the secretion of the Group III substrate, EspE, and the Group IV substrate, EspA, switch in response to acidic pH.

**Figure 4.**
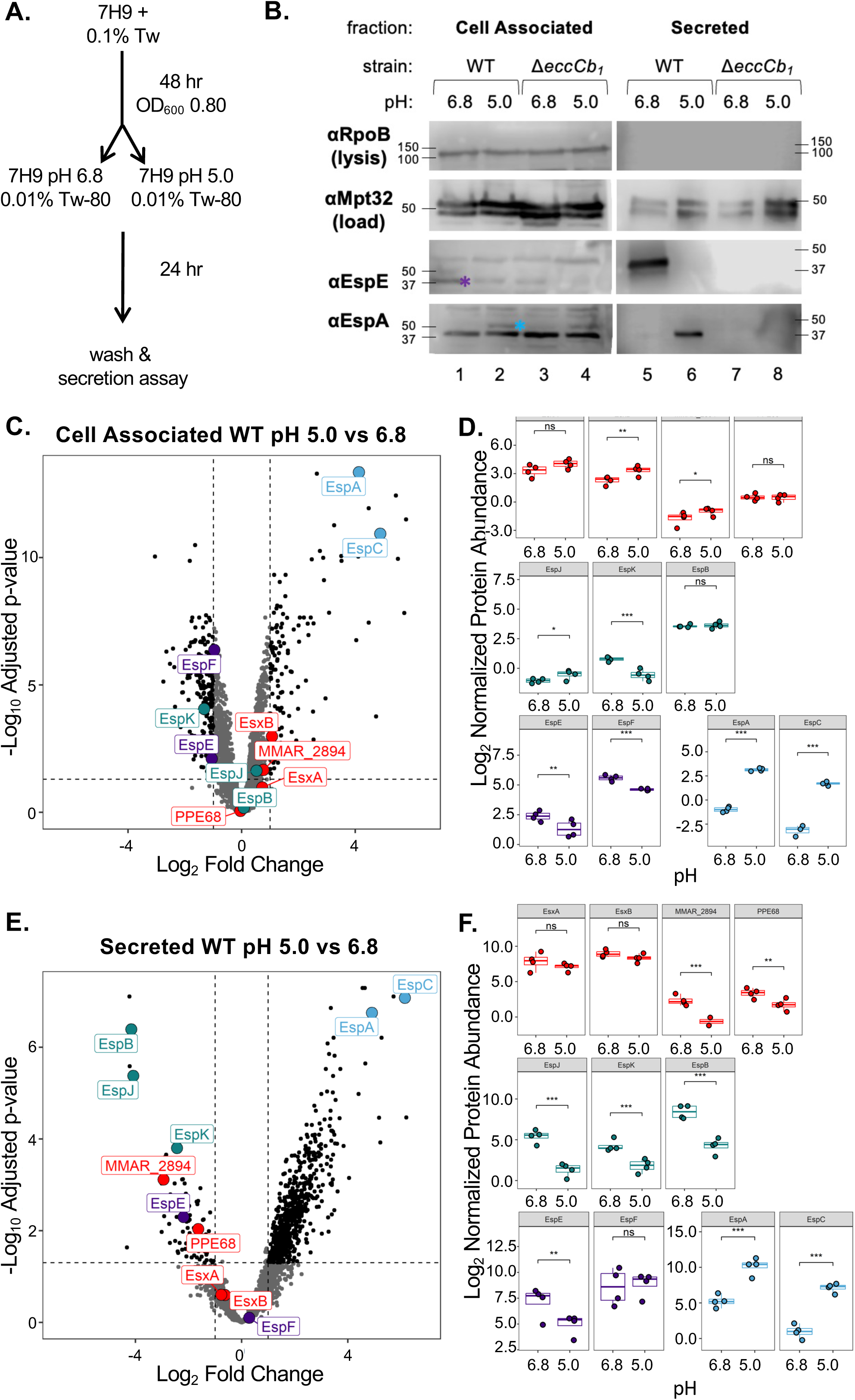
Growth at acidic pH switches ESX-1 substrate secretion. **(A)** Schematic of growth conditions for panel B. **(B)** Immunoblot of 20 µg of *M. marinum* cell-associated (left) and secreted protein fractions (right). RpoB is a control for lysis. MPT-32 is a loading control for the secreted fractions. The blot is representative of three independent biological replicates. Molecular weight markers shown in kilodaltons. The cell associated proteins were resolved on a 12% acrylamide gel. The secreted proteins were resolved on a 4-20% min-protean TGX gel. **(C, E)** Volcano plots show Log_2_ Fold Change (LFC) of WT strains between pH 6.8 and pH 5.0 plotted against the –Log_10_ adjusted p-value for each protein. Dotted lines are drawn at LFC of -1 and 1, and a significance cutoff of adjusted p-value = 0.05. **(D, F)** Box and whisker plots show independent biological replicates (each is an average of two technical replicates) for each protein with the significance annotation from the Benjamini-Hochberg adjusted p-value (ns = not significant, * = adj.p.value < 0.05, ** = adj.p.value < 0.01, *** = adj.p.value < 0.001) as calculated using the biological replicates (n=4).

To determine how growth in acidic pH alters the *M. marinum* proteome, we measured changes to protein abundance and secretion using mass spectrometry-based proteomics. Data Independent Acquisition mass spectrometry was performed (Dataset S1). In the cell-associated fraction of the WT strain we quantified 3215 and 3236 proteins from samples grown at pH 5.0 and pH 6.8 respectively (Figure 4C). We measured significant changes in the levels of cell-associated proteins, with 119 significantly increased and 189 significantly decreased in pH 5.0 over pH 6.8. The levels of EspA and EspC (Group IV substrates) were increased 17- and 29-fold (or 4.12 and 4.87 log_2_ fold change, LFC) at pH 5 compared to pH 6.8. We measured smaller, but significant changes in the levels of the EspK (Group II) and EspE/EspF (Group III) substrates (Fig. 4E). Notably, we did not measure significant changes to the levels of the ESX-1 membrane complex components (EccB_1_, EccCa_1_, EccCb_1_, EccD_1_, EccE_1_ or MycP3, Fig. S6), supporting that the ESX-1 system is functional under both conditions. We measured significant reductions in the cell-associated levels of EccA and EspG, which are proposed to interact with and function as cytoplasmic chaperones for subsets of the ESX-1 substrates (EspF/EspC (52), and PPE68/MMAR_2894 (53, 54), respectively, Fig. S6). We also measured significant increases in the levels of EspD, which is co-transcribed with *espAC* and EspR, a transcription factor regulating the *espACD* operon (55).

In the secreted fraction, we measured 2793 proteins and 2170 proteins from samples grown at pH 5.0 and pH 6.8 respectively (Fig. 4E). Consistent with our prior report (33), the secretion of the ESX-1 substrates was significantly reduced in the Δ*eccCb_1_* strain compared to the WT and Δ*eccCb_1_*/p*eccCb_1_* complemented strains at pH 5.0 and pH 6.8 (Fig. S7 and Dataset S1). Interestingly, we measured a significant increase (30 and 71-fold or 4.90 and 6.15 LFC) in the secretion of EspA and EspC (Group IV substrates) at pH 5.0 relative to pH 6.8. We also measured significant reductions in the secretion of Group I substrates, MMAR_2894 (LFC -2.94) and PPE68 (-1.64 LFC), the Group II substrates (EspJ (-4.09 LFC), EspB (-4.16 LFC) and EspK (-2.45 LFC), and the Group III substrate, EspE (-2.20 LFC), all of which are higher magnitude decreases in pH 5.0 than their LFC in the cell-associated fraction. Together, these data support that the abundance of the Group IV substrates and the ESX-1 chaperones are significantly changed in response to pH. Moreover, the data independently demonstrate pH-dependent substrate switching between the Group II/III and the Group IV substrates.

### Acid stress and infection models reveal changes to ESX-1 transcription and substrate requirements

In Type III secretions systems, substrate switching is regulated through changes in substrate gene transcription (56–58). We performed relative RT-qPCR to measure the transcript levels of ESX-1 substrate genes in *M. marinum* strains grown at pH 6.8 or pH 5.0. We normalized the transcript levels following growth at pH 5.0 to the transcript levels following growth at pH 6.8 (Fig. 5A). The relative levels of the *eccCb_1_* transcript, as well as the *espJ, espK* (Group II, teal) and *espF* (Group IV, purple) transcripts were not significantly different following growth at pH 5.0, compared to growth at pH 6.8. However, the relative levels of the Group I transcripts (red), as well as *espB* (Group II, teal) and *espE* (Group IV, purple) transcripts were significantly reduced following growth at pH 5.0, compared to growth at pH 6.8. Conversely, the transcripts from the Group IV substrate genes, *espA* and *espC,* were significantly increased following growth at pH 5.0 relative to growth at pH 6.8. These data suggest that substrate switching is mediated, at least in part, by the increased transcription or transcript stability of the Group IV substrate genes during growth at acidic pH.

**Figure 5.**
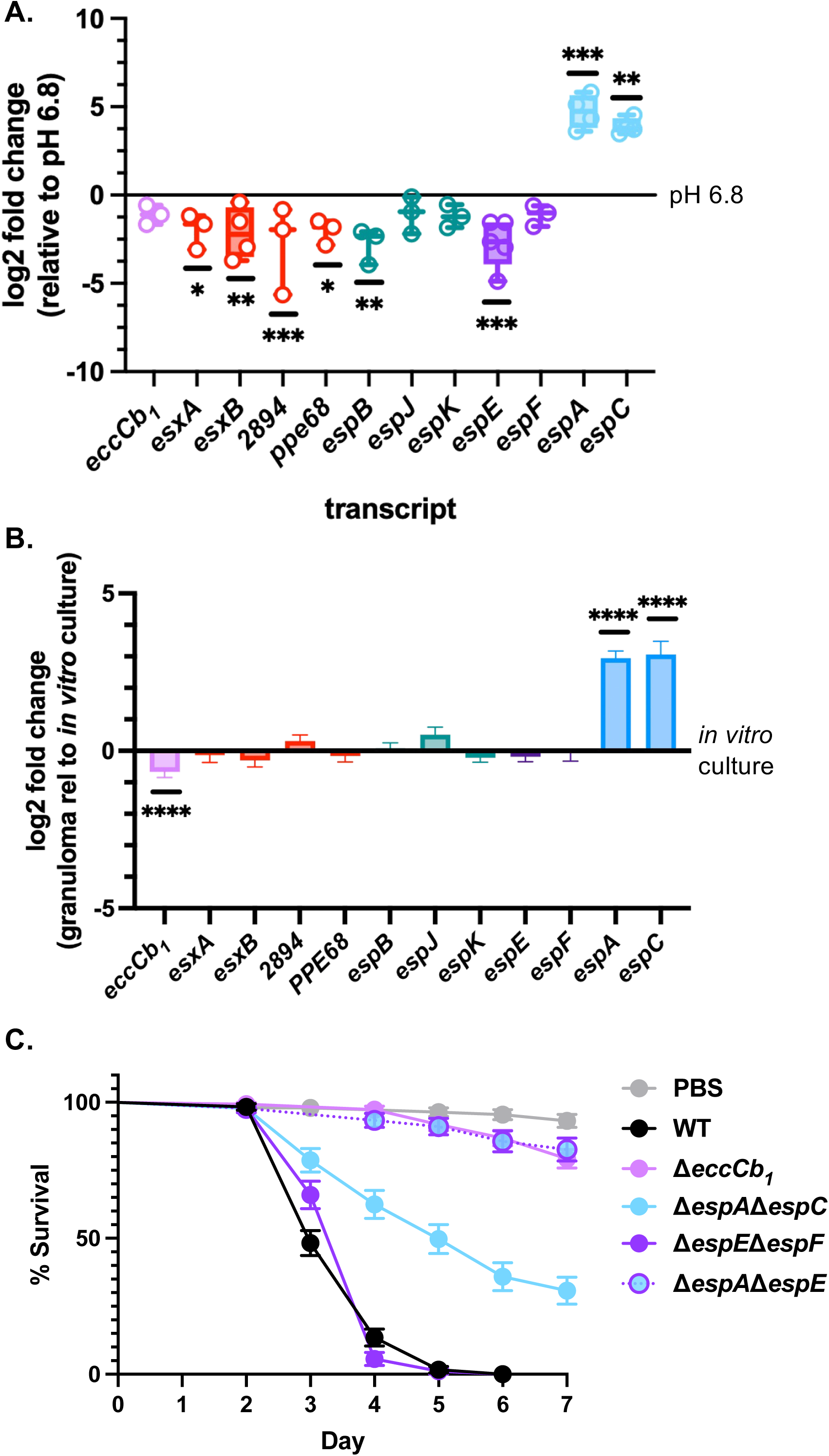
Acid stress and infection models reveal changes to ESX-1 transcription and substrate requirements during infection. (**A)** Log2 fold change of ESX-1 transcript levels in the WT strain following growth at pH 5.0 normalized to transcript levels following growth at pH 6.8. All transcripts are normalized to *sigA* levels. Each data point is a biological replicate, the average of three technical replicates. Statistical analysis was performed using one-way ordinary ANOVA (*P*<0.0001), followed by a Dunnett’s multiple comparison test. Significance is shown relative to the same transcript measured at pH 6.8. *esxA*\**P*=0.0135, *esxB* ** *P*=0.0036, *MMAR_2894* *** *P*=0.0009, *ppe68* * *P*=0.0108, *espB* ** *P*=0.0010, *espE* *** *P*=0.0002, *espA* *** *P*=0.0005, *espC* ** *P*=0.0083. **(B)** RNA-expression analysis of *M. marinum* genes within adult zebrafish granulomas relative to *in vitro* growth, analyzed from RNA-seq data from(92). n=4 independent biological replicates of ∼400 granulomas from 8-12 WT zebrafish for each replicate infected with ∼350 CFU for 14 days and n=3 independent biological replicates for *in vitro* growth. *eccCb_1_* **** adj*P*= 0.0004, *espA* **** adj*P*= 8.32e^-36^, *espC* **** adjP = 5.35e-^12^. (**C**) Percent survival of *G. mellonella* infected with *M. marinum*. Larvae were injected with 10^7^ bacteria in the hindmost proleg, in 3 groups of 10 per biological replicate. Data includes three biological replicates. Statistical analysis was performed using a Log Rank Mantel-Cox Test (*P*<0.0001).

We next tested if ESX-1 substrate switching in response to acid stress was relevant during infection using infection models that are known to have acidic environments. Zebrafish infection with *M. marinum* readily exposes the bacteria to acidic compartments (59). We leveraged the zebrafish model of *M. marinum* infection to test if transcription of the Group III and Group IV ESX-1 substrate genes were differentially upregulated in the granuloma. As shown in Fig. 5B, the *espA* and *espC* transcripts, but not the *espE* and *espF* transcripts, were significantly upregulated in the granuloma compared to the same transcripts following *M. marinum* growth in liquid media. These data support that the Group IV transcripts are significantly increased *in vivo* in a hallmark immune structure of mycobacterial infection associated with lower pH.

Our previously published data demonstrated that in naïve macrophages, the Group III substrates are required, and the Group IV substrates are dispensable for infection by *M. marinum* (33, 37). Our hemolysis data support the hypothesis that the Group III and Group IV both contribute to virulence during acid stress. *Galleria mellonella* larvae are an established, acidic infection model for pathogenic mycobacteria, including *M. marinum* (60–62). The ESX-1 secretion system is essential for killing of *G. mellonella* by *M. marinum* (61, 63). As shown in Fig. 5C, *G. mellonella* infected with the WT *M. marinum* strain (black) had a median survival of 3 days post infection. The median survival of *G. mellonella* injected with phosphate buffered saline (PBS, grey), or infected with the ESX-1 deficient Δ*eccCb_1_* strain (light purple) had an undefined median survival because by the end of the 7-day assay, 93.20% and 79.24% of larvae survived, respectively. Similar to the larvae infected with the WT strain, larvae infected with the Δ*espE*Δ*espF M. marinum* strain, which lacks the Group III substrates, had a median survival of 4 days. The larvae infected with the Δ*espA*Δ*espC M. marinum* strain, which lacks the Group IV substrates, had a median survival of 5 days. These data suggest both the Group III and Group IV substrates can promote *G. mellonella* killing. We hypothesized that the Group IV (EspA and EspC) substrates promoted killing during infection by the Δ*espE*Δ*espF* strain, while Group III substrates (EspE and EspF) promoted killing during infection by the Δ*espA*Δ*espC* strain. To test this hypothesis, we infected *G. mellonella* with the Δ*espA*Δ*espE* strain, which lacks the Group III and Group IV substrates. The *G. mellonella* infected with the Δ*espA*Δ*espE M. marinum* strain had an undefined median survival, with 71.65% of larvae surviving at the end of the 8-day experiment, similar to the Δ*eccCb_1_* strain. Together, these data support that both the Group III and Group IV substrates contribute to virulence in an acidic infection model.

## Discussion

The data presented here directly support that the ESX-1 secretion system in *M. marinum* switches substrate secretion in response to acid stress, both in the laboratory and during infection. In other types of bacterial secretion systems, substrate switching occurs between secreted translocon components, and from the translocon to the secreted effectors (34, 45, 57, 64). Because ESX-1 substrates are secreted in a hierarchy (33), there must be mechanisms and signals that dictate switching between the substrate groups. It remains unclear which substrates are translocon subunits and which are effectors. The Group III and Group IV substrates are encoded from paralogous loci and are dispensable for the secretion of other ESX-1-dependent substrates (33, 37), arguing against their role as part of the translocon. One explanation consistent with our findings is that we are measuring switching between two sets of paralogous effectors that promote lysis of phagosomes or compartments of varying acidity. The roles of specific effectors of conserved secretion systems vary by species. Indeed, there are differences in the literature regarding the importance of the Group IV substrates for virulence in *M. marinum* and in *M. tuberculosis*. Deletion of the *espACD* genes attenuates *M. tuberculosis* and blocks the secretion of the EsxA and EsxB substrates under standard secretion conditions (16, 55, 65, 66). However, deletion of the *espACD* genes does not impact ESX-1 secretion under standard laboratory conditions or the virulence of *M. marinum* during macrophage infection (33). Similar to *M. marinum,* deletion of the *espEF* genes attenuates *M. tuberculosis* in infection models (67, 68). If the Group III and Group IV substrates represent paralogous effectors that function in distinct infection environments, this may explain the reported differences between the two pathogens. *M. tuberculosis* may use a different, hybrid assembly of the ESX-1 system that simultaneously requires the Group III and Group IV substrates. Alternatively, the Group IV substrates may play an expanded role in *M. tuberculosis,* while the Group III substrates have an expanded role in *M. marinum* reflecting their distinct infection niches. Further studies are underway to distinguish these possibilities.

The ESX-1 systems of both *M. marinum* and *M. tuberculosis* lyse the phagosomal membrane. Differences in these secreted effectors could explain the distinct hemolytic activities between *M. marinum* and *M. tuberculosis*. While there are some reports that *M. tuberculosis* has hemolytic activity (35), there are very few compared to those reporting the robust hemolytic activity of *M. marinum.* We were able to exploit the use of *M. marinum* to demonstrate the change in substrate requirement that would not have been obvious when studying ESX-1-dependent phagosomal lysis in an asynchronous infection model.

Our data supports that the Group II substrates, EspB, J and K, are required for the secretion of the Group III substrates, but not the Group IV substrates. Likewise, they are dispensable for lytic activity at acidic pH. It is possible that there are additional, as of yet unidentified substrates akin to the Group II substrates that are specifically secreted under acidic conditions. These groups of substrates may serve as adaptors between the Group I substrates and the Group III/IV substrates. Although numerous studies have identified ESX substrates under standard secretion conditions, our data support that secretion may change in response to the specific signals.

The Group I substrates may be both components of the translocon, because they are required for the secretion of the Group II/III and IV substrates, and effectors in the host. There are also paralogous loci of Group I substrates with no known function. Studies by Shah and Briken suggested that *M. tuberculosis* uses paralogous regions act as accessory ESX-5 systems to secrete distinct subsets of substrates (69, 70). These alternate substrates, which would correspond to Group I substrates here, could be alternate ESX-5 assemblies using three paralogous loci in response to undefined signals. These studies support that ESX systems are dynamic and tailor secretion to the environment.

Our findings indicate that transcriptional regulation likely contributes to substrate switching both during growth in the laboratory and in the zebrafish granuloma infection model. We measured a significant increase in EspR levels following growth at pH 5.0. EspR positively regulates the *espACD* operon in *M. tuberculosis* (55). EspR is regulated by PhoPR, which is a pH responsive two-component system (71). Increased levels of EspA and EspC in the cytoplasm could drive increased secretion following growth under acidic conditions, as supported by our proteomic analyses. There are similar transcriptional connections in substrate switching in Type III secretion systems. However, we think this unlikely because the *espE* and *espF* transcripts remain high in in the zebrafish granuloma. Our prior proteomic analyses suggest competition between the Group II/III substrates and Group IV substrates for secretion *in vitro* (33). Deletion of individual Group II substrate genes, *espB*, *espJ* or *espK*, resulted in significantly increased secretion of the Group IV substrates, EspA or EspC, as measured by proteomics. This finding is consistent with the proteomics data in this study. Indeed, acid stress led to a significant reduction in the secretion of the Group II substrates and a significant increase in the Group IV substrates. In Type III secretion, competition for chaperones or machinery (sorting complex) drives substrate switching in response to environmental signals. The ESX-1 system has two chaperones, EccA and EspG. EspG directly interacts with PE/PPE proteins including PPE68 (53, 54, 72). We previously showed that both EspC and EspF directly interact with the same cytoplasmic ESX-1 component, EccA (52), supporting a potential mechanism for competition. EccA may be part of a sorting complex in the cytoplasm that dictates the order of substrate delivery to the secretion machinery. Interestingly, our proteomics analysis suggested that following exposure to acid stress, the levels of EccA was significantly reduced. It is possible that reduced EccA, with increased EspC/EspA could result in an increase in Group IV secretion and a corresponding decrease in Group III secretion. We also found a significant reduction in EspG levels following acid stress. EspG interacts with PPE68. Reduced EspG could explain the reduction in PPE68/MMAR_2894 secretion measured following growth at pH 5.0. We are continuing to define the mechanisms underlying the initial report of substrate switching observed here.

Mycobacteria encounter environments of varying acidity during infection. Up to 90% of cells infected during mycobacterial infection are macrophages (73). The infected macrophage populations are functionally heterogeneous, controlling or promoting mycobacterial growth (73, 74). Phagosome physiology is likewise heterogeneous. When infecting naïve, resting macrophages, pathogenic mycobacteria arrest phagosome maturation, resulting in a mildly acidic phagosome (pH ∼6.2). However, in activated macrophages, the lysosome fuses with the mycobacteria-containing phagosome, resulting in an acidic phagolysosome (pH 4.5-5) (75–80). Later during infection, mycobacteria are contained within heterogeneous granulomas with a median pH of 5.5 (81, 82). Our findings may inform how pathogenic mycobacteria lyse phagosomes of differing pH, and suggest that pathogenic mycobacteria have the potential to change the virulence factors that mediate phagosomal escape in response to their environment.

## Materials and Methods

Bacterial strains were derived from the *M. marinum* M strain (ATCC BAA-535). Strains were maintained in 7H9 media, and exposed to acid stress using 7H9 buffered with 3-(*N*-morpholino)propanesulfonic acid (MOPS) to either pH 5.0 or pH 6.8 with 0.1% Tween-80 or 0.2% tyloxapol as indicated. *M. marinum* deletion strains were generated using allelic exchange as (33). Hemolytic activity was measured similar to (33, 38), except that strains were grown in buffered and/or detergent treated media for 24 hours and washed prior to hemolysis assay. RNA extraction and RT-qPCR was performed as (83). Protein secretion assays were performed following 48 hours of growth in MOPs buffered 7H9 media (pH 5.0 or pH 6.8) with 0.01% Tween-80 with 0.5% glycerol, but without glucose. Cell associated and secreted protein fractions were generated as in (38). Mass Spectrometry and data analysis were performed as (84). The Protein fractions were precipitated with acetone (85) followed by SDS-PAGE and immunoblot analysis (38). Wax worm infections were performed as (63). An expanded methods section is available in the Supplementary Materials.

## Acknowledgments

This publication was supported by the Institute of Allergy and Infectious Disease of the National Institutes of Health under award numbers R21AI142127, R21AI181133, R01AI106872 and R01AI188782 to PAC, and R01AI130236 to DMT. The content is solely the responsibility of the authors and does not necessarily represent the official views of the National Institutes of Health. We acknowledge all members of the Champion lab for their helpful feedback and discussion. Raw data files, search parameters, and instrument method parameters were deposited in massIVE (identifier MSV000100192) and Proteome Xchange (identifier PXD071871). *Reviewer access at* ftp://MSV000100192@massive-ftp.ucsd.edu*, password: champiomics! *Remove before final publication\**

## Author Contributions

Conceptualization: OAC, RLH, PAC, Methodology: OAC, RJP, CM, GV, RMC, RLC, DMT, MMC, SDW, PAC, Investigation: OAC, RJP, CM, GV, RMC, RLH, Visualization: OAC, RLH, PAC, SDW Funding acquisition: PAC, DMT, Project administration: PAC, Supervision: DMT, MMC, PAC, Writing – original draft: OAC, PAC, Writing – review & editing: OAC, RJP, CM, RMC, GV, RLH, SDW, MMC.

